# Major flaws in “Identification of individuals by trait prediction using whole-genome sequencing data”

**DOI:** 10.1101/185330

**Authors:** Yaniv Erlich

## Abstract

Genetic privacy is an area of active research. While it is important to identify new risks, it is equally crucial to supply policymakers with accurate information based on scientific evidence. Recently, Lippert et al. (PNAS, 2017) investigated the status of genetic privacy using trait-predictions from whole genome sequencing. The authors sequenced a cohort of about 1000 individuals and collected a range of demographic, visible, and digital traits such as age, sex, height, face morphology, and a voice signature. They attempted to use the genetic features in order to predict those traits and re-identify the individuals from small pool using the trait predictions. Here, I report major flaws in the Lippert et al. manuscript. In short, the authors’ technique performs similarly to a simple baseline procedure, does not utilize the power of whole genome markers, uses technically wrong metrics, and finally does not really identify anyone.

## Analysis

### 1. The Venter method is not much better than a baseline re-identification

The results of the authors are unremarkable. I achieved a similar re-identification accuracy with the Venter cohort in 10 minutes of work without fancy face morphology or digital signatures of voices. Instead, I used a simple re-identification procedure that relies on basic demographic information: age, sex, and self-reported ethnicity. These pieces of data are not considered protected identifiers by HIPAA Safe Harbor, and are available in many genomic databases, such as Coriell Cell Repository or the Personal Genome Project (PGP).

To prove that my simple re-identification works for the Venter cohort, let’s examine the demographic data reported in his paper. **Table 1** shows the sex and ethnic background of the samples and was taken directly from Figure 1A of the Venter Study. **Table 2** is an approximation of the age distribution for the samples according to Figure 1C of the Venter study. The resolution of the age distribution is only in ~7-year categories since this is the level of resolution reported in Figure 1C in the Venter paper. I inferred the number of individuals in each age category by measuring the height of the column in the figure. I also repeated the analysis with slightly different numbers and got similar results. Importantly, in real scenarios (e.g. Coriell or PGP) age is usually given at a single year resolution. This means that our analysis is rather conservative because it uses lower resolution data that is expected to be available for the adversary. For the joint age-sex-ethnicity distribution, I assumed that there is no covariance between the age and the sex-ethnic group distribution. Finally, I created a simple R script available on <u>GitHub</u> (https://github.com/TeamErlich/venter_response) to sample from this joint distribution a group of people and test their identifiability.

**Figure 1:**
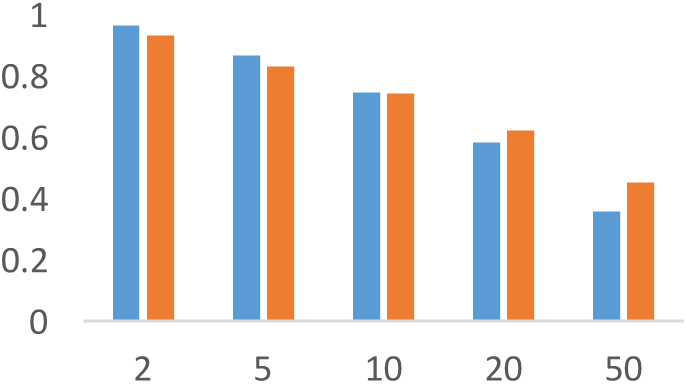
Comparing Venter re-identification to a simple baseline procedure for various n. Blue: results based on the R script with 1000 rounds. Red: Table 2 of Venter (“Full-Select” row).

**Table 1:** sex and self-reported ethnicity of the Venter cohort

|  | African | European | Latino | East Asian | South Asian | Other |
| --- | --- | --- | --- | --- | --- | --- |
| Female | 381 | 126 | 39 | 39 | 8 | 39 |
| Male | 188 | 147 | 24 | 24 | 10 | 36 |

**Table 2:** Approximation of the age distribution of the Venter cohort.

| Age Group | #1 | #2 | #3 | #4 | #5 | #6 | #7 | #8 | #9 | #10 |
| --- | --- | --- | --- | --- | --- | --- | --- | --- | --- | --- |
|  | 295 | 190 | 130 | 90 | 95 | 120 | 100 | 35 | 5 | 1 |

The Venter study defined success as having a unique combination of identifiers for the sample of interest within a group of *n* individuals. It is worth saying that this is a very weak definition of success for various reasons outlined below. But for now, we will use the same definition for compatibility with their study and set n=10 because this is the value reported in the PNAS abstract.

Venter reported 74% success rate after employing close to ~30 authors and a range of complex algorithms. To contrast, the simple baseline demographic re-identification procedure got 75% success rate after one hour of work. I also tested other group sizes, namely n=2, 5, 10, 20, and 50. In nearly all conditions, the simple procedure was comparable to the Venter method (**Figure 1**). The only exception was 50 individuals, however, if I had access to a single year age resolution, similar to a real scenario, it is very likely that the baseline procedure would have had higher success rates also to this condition.

The take home message should be that identifying someone in a group of ten people requires very little effort. Anyone with access to even low dimensional data, such as basic demographic, can do that. This is not very surprising. It takes only log(10)/log(2)=3.34bits of information to reduce 10 possibilities into one option. In our previous work, Arvind Narayanan and myself (Nature Rev. Genetics, 2014; Box 1) assessed the information content of simple demographic identifiers such as age or state of birth. Many of these identifiers give >5 bits of information, meaning that each one of them is likely to be unique in a group of 10 people.

To summarize, Venter’s team had to use a very poor success criterion in order to show that the accuracy is high (the abstract reported 80%). But effectively, much simpler techniques could achieve a similar success rate.

### 2. No face value

The claimed novelty and impact of the Venter manuscript is using trait predictions to identify people (especially face predictions). Unfortunately, careful examination of the figures shows that the re-identification power virtually stems from inferring genomic ancestry and sex from the genetic data rather than trait-specific markers. Thus, the authors do not infer the height or the face structure of a specific person. Instead, they infer something that is very close to the population average and use this value to re-identify the sample. It is like the authors needed to get to the supermarket and decided to build a spacecraft to make the trip.

For example, the authors reported a R^2^=0.53 for height prediction. I was first highly surprised by this measure since a GWAS study of quarter million people — 200x the size of the Venter dataset — explained only 16% of the height variance (Wood et al., Nature Genetics, 2014). However, a more careful look at Figure 5B (see below) shows that nearly 80% of the explained height variance of the Venter predictions (that is R^2^=0.4 out of R^2^=0.53) is due to the inclusion of sex markers! This means that the authors do not really predict personal height. Rather, they mainly infer sex averages of height. To make things worse, the other 10% of the explained height variance was driven by adding ancestry markers. Again, the authors do not really predict individual level height from genetic data (or at least this is not what gives the re-identification power). Rather, they infer very simple demographic traits and use those traits to find population averages rather than a personal prediction.

**Figure 5B.**
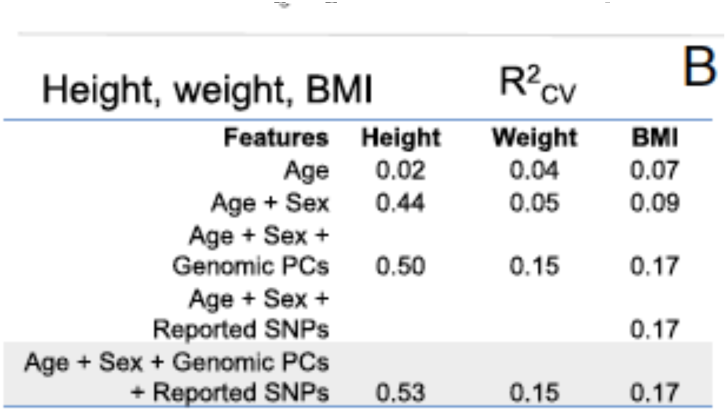
from the Venter study.

Face prediction works poorly as well. Figure 3 of Venter shows that sex and genomic ancestry are the most important covariates for face morphology. Adding the associated SNPs barely improves the prediction. This is not surprising. Face is a complex, high dimensional trait with age-by-genetic interactions. The field cannot accurately predict height, a scalar trait that barely changes during most of adulthood despite GWAS of 250,000 people. So it is obvious that it does not work.

**Figure 3.**
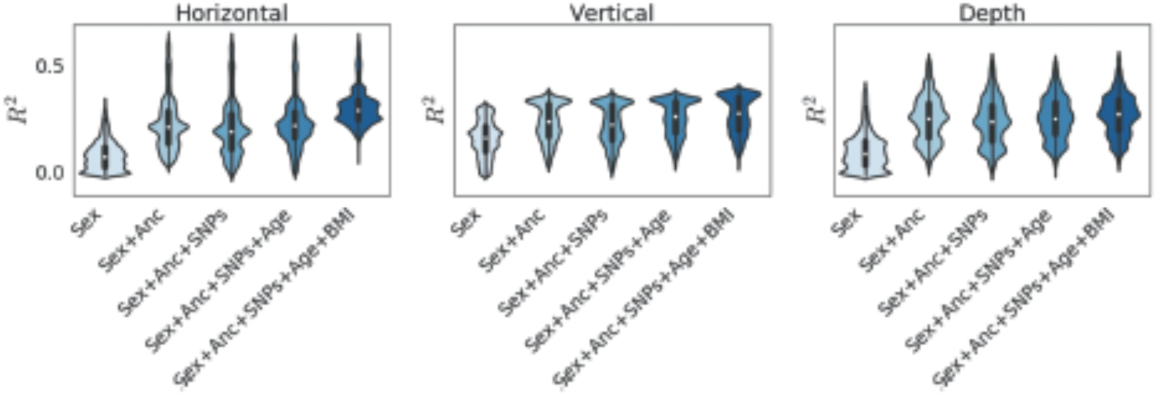
from the Venter study.

Figure S11 in the Venter study shows that population means are predicted rather than actual individual level data (courtesy for Jason Piper for the idea). The left column is the true face of the person and the right column is the predicted face:

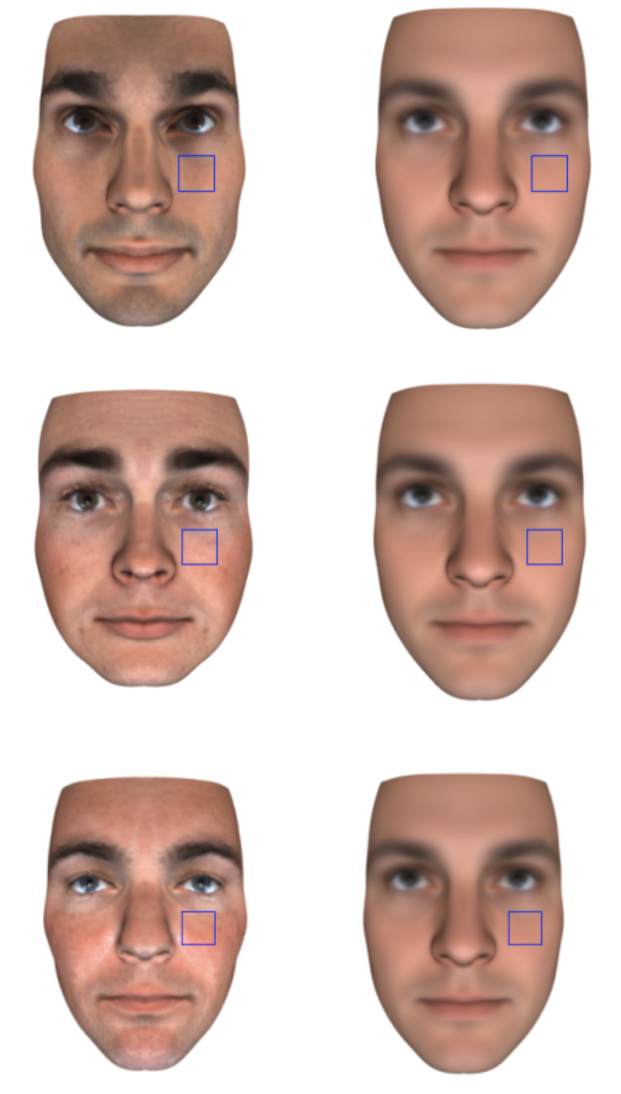
Examples from figure S11 from the Venter study

For each of these three different white males, the predicted faces are virtually identical. Clearly, the Venter algorithm predicts a “genetic white male face” rather than the individual level face of the person.

To summarize this point, the title says: “Identification of individuals by trait prediction using whole-genome sequencing data” but most of the trait predictions is carried by ethnicity of the individual (genomic PCs) rather than the trait specific SNPs. Also do not by that excited about the 1000 genomic PCs. The first author (which is honest about the issues in the study) acknowledged on Twitter that the high PCs (>100) explains virtually nothing of the ethnicity variance.

### 3. Technical sensitivity

One nice contribution of the Venter study is inferring age from whole genome samples. However, this method is unlikely to work in order to breach sample anonymity in the real world. The authors indicated in multiple locations that their ability to predict traits hinges on the consistency of their pipeline. Specifically, for age, the authors stated that their age prediction worked only after sequencing NA12878 for 512 times using the same exact sequencing pipeline. Consider an adversary that wants to use the Venter technique and predict the age of a random dbGAP sample. It means that the adversary needs somehow to gain access to the sequencing pipeline and sequence the same sample hundreds of times using the exact parameters. Good luck with that.

Previous studies reported that prediction of highly-dimensional traits from DNA can be very sensitive to the phenotypic capture platform. For example, Schadt et al. (Nature Genetics, 2012) explored a similar problem to the current study: re-identifying genomic samples when the observed traits are gene expression profiles. The Schadt method worked extremely well when the training set and the test set were done using the same array manufacturer. However, they reported poor performance when the training set and test set were done on different array companies. I am highly concerned that this is also the case with the Venter study, especially for face morphology, RGB colors of the eyes, and skin tone.

### 4. The complete registry fallacy

Recall that Venter success criteria was determining whether the person of interest has a combination of unique attributes in a sample of a small number of people. Indeed, the privacy literature usually use the uniqueness of the attributes in a dataset as surrogate for identifiability (for example, *k*-anonymization). However, this measure is used in the context of evaluating <u>anonymization algorithms</u> under the worst case scenario of a powerful adversary with access to a complete registry of the population. Importantly, the Venter team did not develop a protection algorithm; rather, they aimed to demonstrate the utility of a new exploit. It is simply wrong to claim that just because as sample is unique, it means that it has been identified.

To work in the real world, an adversary using the Venter technique would have to create population-scale database that includes height, face morphology, digital voice signatures and demographic data of every person they want to identify. While the proliferation of social media data made it easy to collect phenotypic measures, the ability to collect, extract, standardize, and query a population scale database like that would be paramount. Moreover, the HLI-funded authors harnessed a sophisticated camera system and highly standardized pipeline to collect faces (“high-resolution three-dimensional (3D) system equipped with nine machine vision cameras and an industrial-grade synchronized flash system; the 3D 200-degree face was captured in approximately 1.5 milliseconds”). The US Justice system still struggles to move from STRs to SNP-based analysis. Therefore, it is again totally unrealistic to assume that such phenotypic data is available for adversaries. Even powerful intelligence agencies cannot collect such data for every person in their target population.

### 5. Did the authors actually re-identify anyone?

Final thought, in our previous study (Gymrek et al., Science, 2013), we demonstrated that we can re-identify individuals in a real scenario (ironically, Venter’s genome was one of the successful test cases). If Venter’s method works so great, why not trying to do the same? Why the authors did not simply pick a person from any of the public research projects with the proper consent (1000Genomes, PGP, OpenSNP) and showed that they can identify the sample?

I think that the answer is clear.

